# Genome-wide CRISPR Screens Identify Ferroptosis as a Novel Therapeutic Vulnerability in Acute Lymphoblastic Leukemia

**DOI:** 10.1101/2022.02.01.478478

**Authors:** Marie-Eve Lalonde, Marc Sasseville, Anne-Marie Gélinas, Jean-Sébastien Milanese, Kathie Béland, Simon Drouin, Elie Haddad, Richard Marcotte

## Abstract

<u>A</u>cute lymphoblastic leukemia (ALL) is the most frequent cancer diagnosed in children. Despite the great progress achieved over the last 40 years, with cure rates now exceeding 85%, refractory or relapsed ALL still exhibit a dismal prognosis. This poor outcome reflects the lack of treatment options specifically targeting relapsed or refractory ALL. To address this gap, we have performed whole-genome CRISPR/Cas drop-out screens on a panel of seven B-ALL cell lines. Our results demonstrate that while there was a significant overlap in gene essentiality between ALL cell lines and other cancer types survival of ALL cell lines was dependent on several unique metabolic pathways, including an exquisite sensitivity to GPX4 depletion and ferroptosis induction and GPX4 KO. Detailed molecular analysis of B-ALL cells suggest that they are primed to undergo ferroptosis as they exhibit high steady-state oxidative stress potential, a low buffering capacity, and a disabled GPX-independent secondary lipid peroxidation detoxification pathway. Finally, we validated the sensitivity of B-ALL to ferroptosis induction using patient-derived B-ALL samples.

## Introduction

Acute lymphoblastic leukemia (ALL) is the most prevalent cancer during childhood, representing nearly 80% of all cancer in this age group^1^. Treatment protocols have greatly improved over the last 30 years, such that the survival rate reaches >85%^2^. Despite this therapeutic success, the prognosis for relapsed patients remains dismal with a less than 50% 5 year survival, still making ALL the second highest cause of death by disease amongst children in Canada and in the U.S (Vrooman and Silverman 2016). In addition, up to 65-70% of pediatric ALL survivors will suffer from long-term debilitating or even life-threatening treatment related sequelae^3,4^. Hence, novel therapeutic avenues that are both more effective at achieving long-term remission while eliciting less acute long-term toxicities than current treatment regimen are required to treat these patients.

Recent advances in the use of clustered regularly-interspaced short palindromic repeat (CRISPR)/Cas9 technology has revolutionized functional genomics and the analysis of gene function in mammalian cells with its precision, ease of use, speed, and versatility. Whole-genome bulk pooled CRISPR screens, using multiple sgRNAs targeting each human gene in a single CRISPR library, have allowed the identification of genes implicated in processes underlying phenotypic read-outs such as proliferation and survival in hundreds of tumoral cell lines^5–7^. These large datasets revealed panels of core essential and pan-cancer genes and unraveled key players required for cell viability/proliferation in cell lines derived from multiple histotypes. Although ALL is a common form of cancer, very few ALL cell lines have been previously screened by large dropout screen studies^5–7^. This is likely explained by the difficulty in infecting pre-B and T lymphocytes with lentiviruses, a technical requirement for performing whole-genome CRISPR screens.

In this report, we performed whole-genome dropout CRISPR screens of seven B-ALL cell lines. These screens revealed a surprisingly large subset of essential genes unique to ALL cell survival that were not reported to be core essential genes in previous studies^6–8^. These B-ALL-enriched genes were implicated in different cellular pathways and functions, many of which were associated directly or indirectly to ferroptosis. In recent years, this non apoptotic pathological cell death has gained increasing attention in cancer research, particularly for its potential tumor-suppressor function that could be exploited for neoplastic disease treatment^9–11^. By using different cellular and molecular biology approaches, we validate that B-ALL cell lines have an exquisite vulnerability to ferroptosis induction. This sensitivity is illustrated by the extreme responsiveness of cells to glutathione peroxidase 4 (GPX4) inhibition, but also to other pathways that regulate GPX4 activity. We also show that this sensitivity is rescued by exogenous expression of FSP1, a recently characterized ferroptosis inhibitor^12,13^ that we found to be poorly expressed, not only in B-ALL cells, but in leukemias in general. Finally, we demonstrate that this acute sensitivity to ferroptosis induction is also observed in a panel of B-ALL patient-derived tumor samples.

## Results

### Whole-genome CRISPR screens of B-ALL cell lines

In order to identify genes that are essential for B-ALL cells, we performed whole-genome pooled dropout CRISPR screens using a two-vector system (Fig. 1A)^14^. First, a stable pool of Cas9-expressing cells was established for each cell line. This preliminary step was limiting for many ALL cell lines, since several pools did not reach the 75% Cas9 activity threshold required for further screening (Suppl. Table 2). For some of these cell lines, Cas9 activity decreased rapidly after blasticidin selection, particularly in T-ALL cell lines (data not shown), suggesting that constitutive Cas9 expression might be toxic, similar to what is reported in AML cell lines^14^. Of all the pools tested, the 7 B-ALL Cas9-stable pools attaining >75% Cas9 activity were infected with a whole-genome lentiviral 90K single-guided RNA (sgRNA) library (Suppl. Fig. 1A and B)^14^. Screen quality was very high, as shown by the BAGEL essential and non-essential genes^8,15^ comparison precision/recall curves (Suppl. Fig. 1C). Furthermore, principal component analysis (PCA) shows that T0 time points from all cell lines are tightly clustered and, despite some variance shown in the final timepoint (Tf) between cell lines, cell line replicates were tightly clustered and significantly different from T0, suggesting that significant dropout was achieved in the Tf samples (Suppl. Fig. 1D).

**Figure 1.**
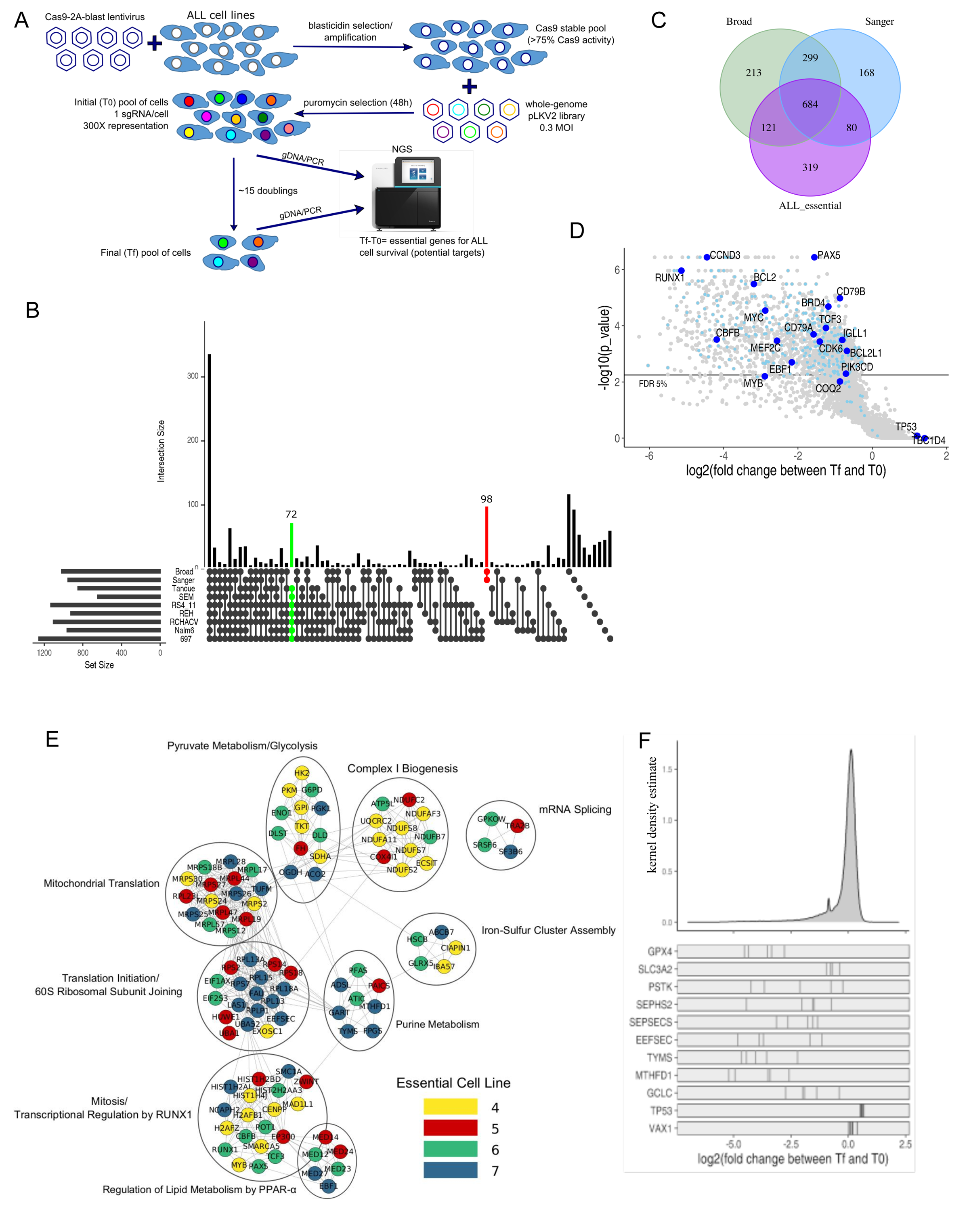
Whole-genome CRISPR/Cas9 screens of ALL cell lines identify ALL essential genes. A) Schema of B-ALL screening pipeline B) Significant genes set intersection of 7 B-ALL cell line CRISPR screen results with UpSet, “Broad” and “Sanger”(Dempster et al. 2019) essential gene list at false discovery rate (FDR) <0,05. The intersections with more than 5 genes are shown. C) Overlap between B-ALL essential (>4/7 cell lines, FDR <0.05) genes, “Broad” and “Sanger” core essential genes. D) Volcano plot of 697 cells CRISPR screen results with FDR <0.05. B-ALL-known dependencies are highlighted in blue and ALL enriched essential genes are highlighted in light blue. Volcano plot of 697 cells is shown as a representative Volcano plot of B-ALL cells screen results. E) Gene Network Analysis of B-ALL enriched essential genes where nodes represent genes and edges represent protein-protein interactions between genes. Node color represents the number of ALL cell lines essential for the given gene. F) Log2 fold change distribution of sgRNAs in the 7 ALL screened cell lines. sgRNAs Log2 fold change for ferroptosis related genes are illustrated at bottom of the panel. VAX1, negative control. TP53 was enriched in several B-ALL cell lines.

### Whole-genome CRISPR screens identify vulnerabilities specific to B-ALL

In an attempt to identify essential gene unique to B-ALL, we compared the essential genes identified in the screened B-ALL cell lines with the list of essential genes reported in previous datasets. We noticed a considerable overlap of B-ALL essential genes with Broad and Sanger core essential genes^7^ (Fig. 1B and C, Suppl. Table 3 and 4). 1204 essential genes were identified in at least 4/7 B-ALL cell lines (FDR < 0.05), of which 67% and 63% were also included in the Broad and Sanger gene lists, respectively and, as expected, belong to core biological processes such as the cell cycle, mRNA translation, splicing, and Pol II transcription-pathways that are vital for growth and survival of most cell lines. Strikingly, 72 genes were identified as essential in every B-ALL cell line screened but were not previously identified as essential genes in the Broad and Sanger datasets (Fig. 1B, green line). Likewise, 98 genes were previously defined as essential in both the Sanger and Broad dataset but were not essential in any of the ALL cell lines (Fig. 1B, red line). We defined as “B-ALL-enriched essential genes” all hits that were found in at least 4/7 of B-ALL cell lines, but that were not present in the Broad or Sanger essential gene datasets (a total of 319 genes as shown in Fig. 1C). These genes are enriched for ALL-associated transcription factors, such as *PAX5, EBF1, CBFB, TCF3*, and *RUNX1*^16–18^, B-cell receptor and signaling, such as *CD79A, CD79B* and *PIK3CD* (Chu and Arber 2001; Kruth et al. 2017; Serafin et al. 2019), and other known B-cell vulnerabilities, such as *CDK6, CCND3*, and *BCL2*^22–24^ (Fig. 1D). A few genes were also enriched in screened ALL cell lines, such as TP53 (most B-ALL cell lines have wild-type TP53) and TBC1D4, indicating that KO of these genes provided a proliferative advantage (Fig. 1D, F, Suppl. Table 3). Mapping the “B-ALL-enriched essential genes” onto the STRING database revealed that these genes were members of specific functional subnetworks with multiple protein-protein interactions (Fig. 1E), such as RUNX1-transcriptional regulation, iron-sulfur cluster assembly, and regulation of lipid metabolism genes. WikiPathway enrichment analysis also identified glutathione metabolism, one carbon metabolism and pentose phosphate pathways, all of which would impact ferroptosis, and ferroptosis regulation itself as specific pathways for B-ALL cell survival (Suppl. Table 5). Ferroptosis is an iron-dependent form of necrosis which is triggered via lipid peroxidation of polyunsaturated fatty acid (PUFA) at the cell membrane^25^ and has garnered significant interest in the past few years as an alternative drug-induced cell death mechanisms to therapy-induced apoptosis-resistant cancers^9^. Central to this cell death pathway is GPX4, one of the top-ranked essential gene according to our screen results (Fig. 1F, suppl. Table 3) and is the main inhibitor of ferroptosis induction. Interestingly, multiple genes potentially impacting GPX4 activity were included in “B-ALL-enriched essential genes” list (Fig. 1F). Hence, our whole-genome screens of B-ALL cell lines identified several genes unique to this histotype and enriched for specific functional pathways, several of which could directly impact ferroptosis induction.

### B-ALL cell are extremely sensitive to ferroptosis induction

GPX4, a selenocysteine-containing glutathione peroxidase, reduces phospholipid hydroperoxides to lipid alcohol with the help of glutathione (GSH) as an obligate cofactor and acts as the main endogenous inhibitor of ferroptosis induction^26^. To validate that inhibition of GPX4 induces ferroptosis in ALL cells, we treated cells with RSL3, a direct inhibitor of GPX4^26^. Comparison of dose-response curves in B-ALL to non-ALL and GPX4 non-essential cell lines (A549, MCF7, NCIH226) indicates that B-ALL cell lines are particularly sensitive to RSL3 treatment (Fig. 2A). This sensitivity was significantly higher compared to the RSL3 sensitivity of many other cell lines reported in previous studies^12,27^. To confirm that RSL3 treatment induce ferroptosis but not apoptosis, we stained ALL cells with C11 BODIPY™, which stains peroxidated lipids, and Annexin V, respectively (Suppl. Fig. 2A). Lipid peroxidation, but not Annexin V staining, was only detected in the RSL3-treated cells (Suppl. Fig. 2A). Notably, higher lipid peroxidation levels were also found in steady-state conditions in B-ALL cells compared to non-ALL cells (Suppl. Fig. 2B). To confirm the sensitivity of ALL to ferroptosis induction genetically, we generated GPX4 knock-out (KO) clones using CRISPR/Cas9 by constantly growing targeted cells in the presence of ferrostatin-1, a radical trapping agent, which strongly inhibits ferroptosis (Fig. 2B and C)^25,27^. Withdrawal of the drug in GPX4 KO clones led to cell death within 18 hours after removal, which confirms the rapid induction of cell death after GPX4 inhibition in B-ALL cells (Fig. 2B). RSL3 drug treatment could partially be rescued by ferrostatin-1, the iron chelator deferoxamine (DFO), as well as PD146176, a 15-lipoxygenase inhibitor, and EUK-134, a general antioxidant (Fig. 2D), but not with Z-VAD-FMK, a pan-inhibitor of caspases and apoptosis. These results indicate that RSL3 creates a lethal oxidative stress environment that is, to some extent, iron-dependent. Interestingly, only lower DFO concentration that was previously reported could rescue cell viability^25,27,28^. This discrepancy can be explained by the fact that B-ALL cells are more sensitive to elevated DFO treatment (Suppl. Fig. 2D), which could reflect their higher requirement for iron for sustained proliferation compared to other normal cell types and malignancies^29^.

**Figure 2.**
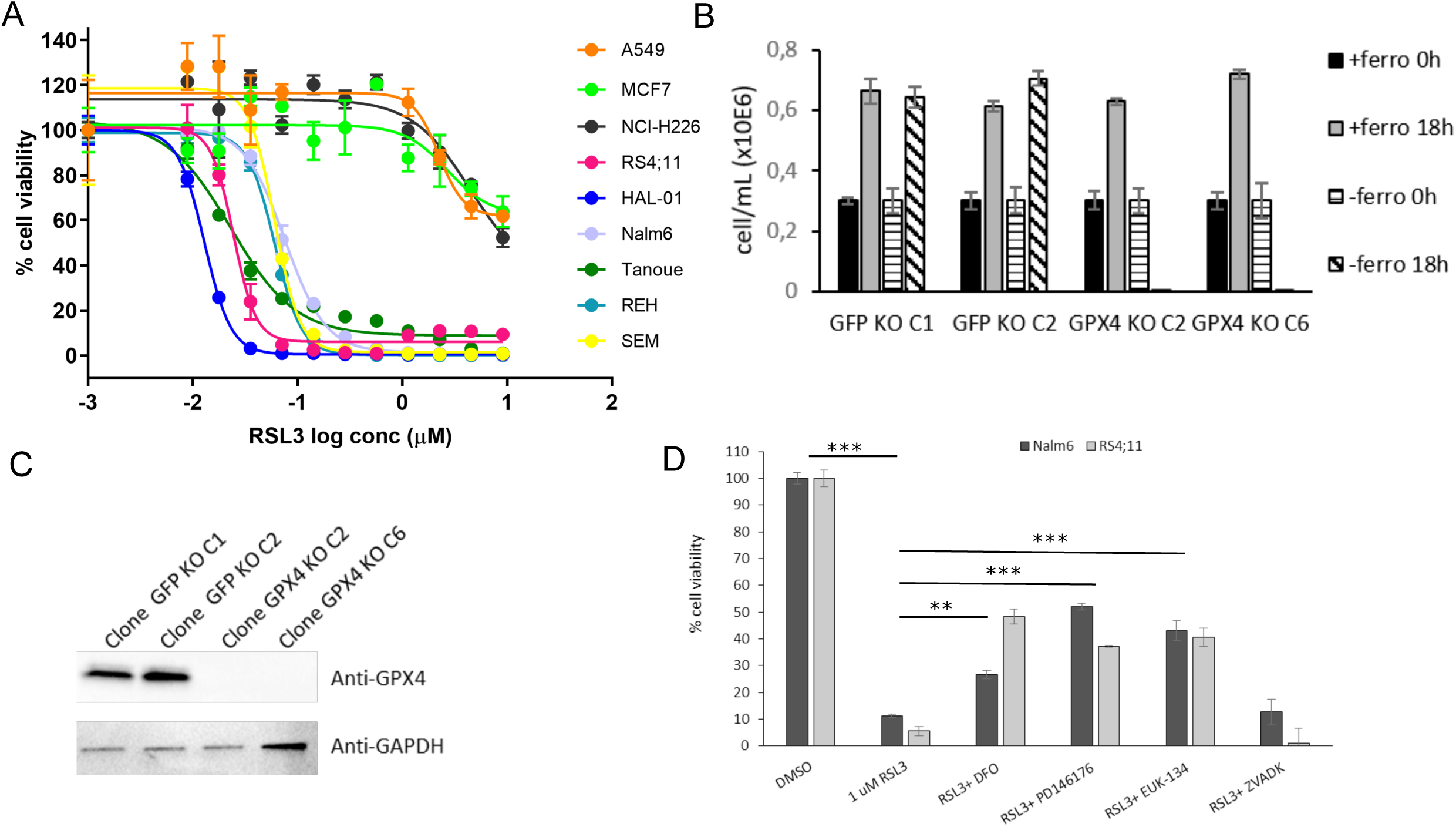
B-ALL are highly sensitive to GPX4 inhibition. A) Cell viability curves of B-ALL and other cell lines (A549, MCF7, NCI-H226) to the GPX4 inhibitor (RSL3). B) Growth of REH GFP clones 1 and 2 (negative control) and REH GPX4 KO clones 2 and 6 +/- ferrostatin (2 µM) for 18h. C) WB analysis of GPX4 protein level in GPX4 KO vs GFP KO clones. D) RSL3 (1 µM) rescue experiments with the iron chelator deferoxamine (DFO; 10 µM)), the pan-caspase apoptotic inhibitor ZVADK (50 µM), the 15-lipoxygenase-1 inhibitor PD146176 (0.5 µM), the superoxide dismutase mimetic EUK-134 (30 µM), or the radical trapping agent ferrostatin (2 µM) in NALM6 and RS4;11 for 24h.**Unpaired t-test pvalue <0,001 and ***pvalue <0,0001.

### B-ALL cells are sensitive to perturbations in pathways regulating GPX4 activity

Because of its requirement for specific cellular metabolites, GPX4 activity and ferroptosis induction are influenced by many metabolic pathways^9^. Several genes regulating these pathways were identified within B-ALL-enriched essential genes (Fig.s 1F and 6). These include genes implicated in the synthesis of GSH (SLC7A11/SLC3A2/GCLC), mevalonate (HMGCR/MVD/MVK), lipids (ASCL3/4, FASN, MCAT), selenocompounds (SEPSEC, EEFSEC, SEPHS2), and iron metabolism (PCPB2, TFRC, STEAP3).

Interestingly, most B-ALL cells showed significantly lower levels of SLC7A11 and GSH compared to non-ALL cells, in steady-state conditions (Fig. 3A and B). Because of its essential role for GPX4 activity and in oxidative stress protection, low GSH levels could contribute to the increased sensitivity of B-ALL to ferroptosis induction. Accordingly, B-ALL cell lines, with the exception of NALM6 whose sensitivity was intermediate, were more sensitive to buthionine sulfoximine (BSO), a glutamate-cysteine ligase (GCLC) inhibitor^30^, and to erastin (a direct inhibitor of the Xc^-^ transporter responsible for L-cystine import), than non-ALL cell lines where GPX4 is non-essential (Fig. 3C, D and E). Rescue of BSO or treatments in NALM6 and RS4;11 cells by addition of exogenous GSH indicates that GSH synthesis is essential for ferroptosis inhibition in B-ALL lines (Fig. 3E, suppl. Fig. 3A).

**Figure 3.**
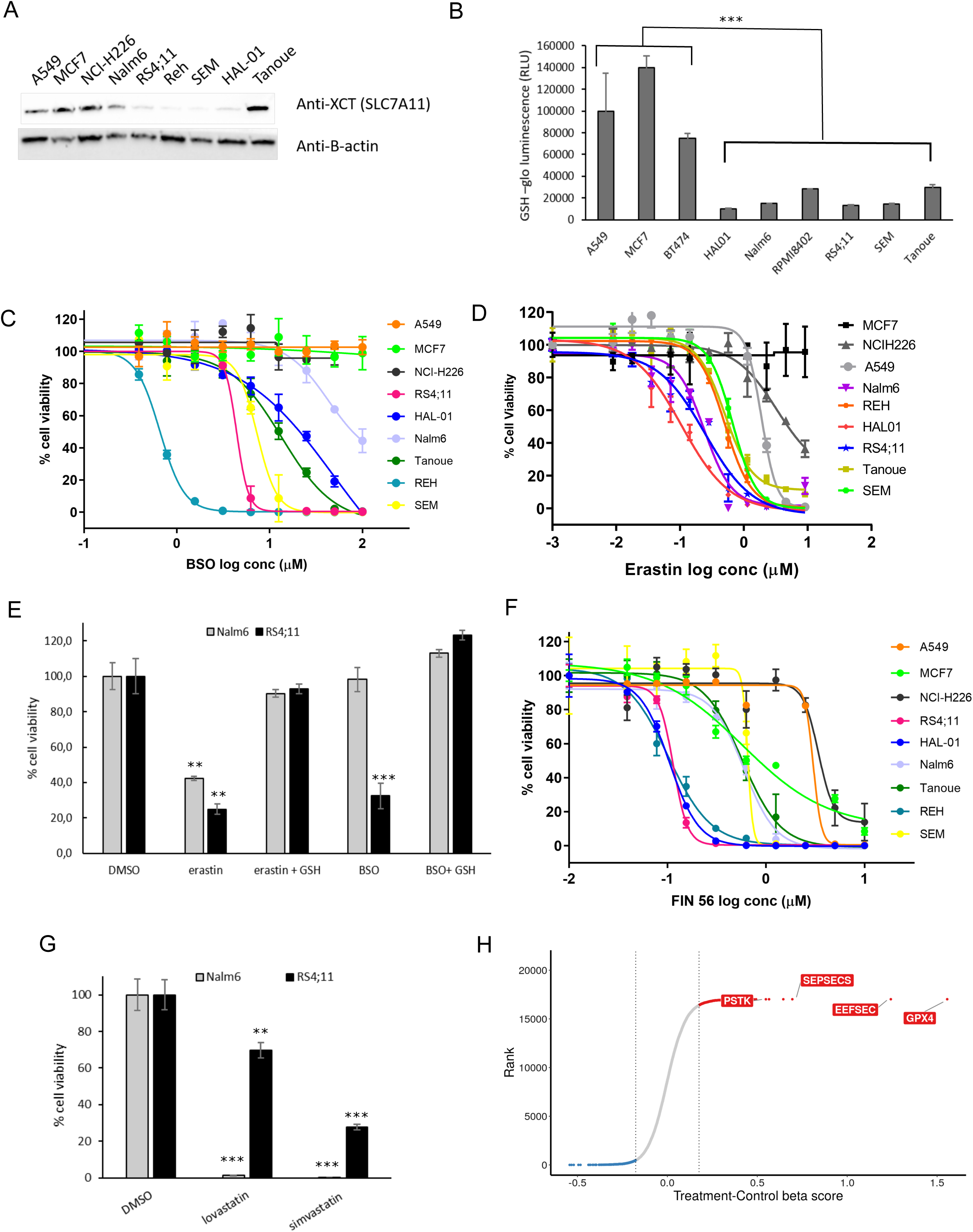
Pathways controlling GPX4 activity contribute to ALL sensitivity. A) WB anti-SLC7A11 in B ALL cell lines. β-actin is used as loading control B) GSH level measurement in steady-state conditions for ALL vs non-GPX4 sensitive (non-ALL) cells. ***Unpaired t-test pvalue <0,0001. C) IC50 curves of BSO in ALL cell lines vs non-GPX4 sensitive (non-ALL) cell lines after 96h treatment. Data shown are from one representative experiment. D) IC50 curves of Erastin inhibitor in ALL vs non-GPX4 sensitive (non-ALL) cells. Data shown are from one representative experiment. E) 10 uM erastin- (system Xc- transporter inhibitor) and 100 uM BSO-treated NALM6 and RS4;11 are rescued by 1mM GSH (48h). *** Unpaired t-test pvalue <0,001 and ** pvalue <0,01. F) IC50 curves of Fin56 inhibitor in ALL vs non-GPX4 sensitive (non-ALL) cells. Data shown are from one representative experiment. G) Cell viability of NALM6 and RS4;11 following treatment with lovastatin (20 uM) and simvastatin (20 uM) for 72h. *** Unpaired t-test pvalue <0,002 and ** pvalue <0,01. H) Beta scores for +/- ferrostatin rescue screens performed in REH^cas9^ and SEM^cas9^ pools. Only GPX4 and selenocompound metabolism genes are indicated. Values represent log2 fold-change between untreated (control) and treated samples.

The mevalonate pathway regulates lipid peroxidation by two different mechanisms. First, it produces CoQ10, which contributes to detoxification of lipid reactive oxygen species (ROS)^31^. Second, it regulates the production of Isopentenyl pyrophosphate (IPP), a metabolite required for Sec-tRNA (a tRNA depositing selenocysteine on proteins) synthesis^31^. Since GPX4 contains a selenocysteine residue essential for ferroptosis inhibition^32^, regulation of selenocysteine incorporation into GPX4 can directly influence its activity. B-ALL cells were more sensitive to inhibition of the mevalonate pathway using FIN56, a dual inhibitor of squalene synthase (an enzyme involved in cholesterol biosynthesis)^33^ and GPX4 in contrast to non-ALL cells (Fig. 3F, Suppl. Fig. 3A). Further confirming our observations, treatments of B-ALL cells with other mevalonate pathway inhibitors (lovastatin, simvastatin) also induced ferroptosis (Fig. 3G).

In an attempt to determine which genes identified as essential in our primary screens dropped out of the pool specifically because of ferroptosis induction, we performed whole-genome CRISPR screens in two Cas9-expressing B-ALL cell lines in the presence or absence of ferrostatin-1 (Suppl. Fig. 3B, Suppl. Table 6). In this setting, we would expect genes depleted in the untreated group because of ferroptosis induction to no longer being depleted in the ferrostatin-treated group. As expected, GPX4 sgRNA were no longer depleted in the presence of ferrostatin-1 and showed the highest fold change between untreated and treated samples (Fig. 3H). Furthermore, three genes implicated in selenocysteine synthesis (PSTK, EEFESEC and SEPSECS) were also in the top hits further arguing that selenocysteine levels regulate ferroptosis in B-ALL cells (Fig. 3G, Suppl. Table 6). Cells in which these genes were individually knocked out using CRISPR/Cas9 also showed increased lipid peroxidation (Suppl. Fig. 3C), strongly supporting ferroptosis induction. These results are consistent with DepMap CRISPR screen data. By segregating cell lines as GPX4 essential or GPX4 non-essential (Suppl. Fig. 3D, see materials and methods) and looking for genes that are co-essential with GPX4, we observed co-essentiality of SEPSECS, SEPHS2, EEFSEC, PSTK and SECISBP2 genes (Suppl. Fig. 3E), which are all selenocompound metabolism genes.

### B-ALL cells express low levels of FSP1, a potent ferroptosis inhibitor

Distribution of GPX4 mRNA expression and protein levels between GPX4 sensitive and non-sensitive cell lines in the DepMap/CCLE indicate that GPX4 expression level does not account for GPX4 essentiality in cells (Suppl. Fig. 4A and B). When looking at differentially expressed genes between these two groups we noticed that FSP1/AIFM2, a recently characterized ferroptosis inhibitor^12,13^, was expressed at lower levels in cells that were dependent on GPX4 compared to non-GPX4 essential cells (Suppl. Fig. 4C). A similar trend was also seen at the protein level (Fig. 4A). Moreover, mRNA and protein levels of FSP1 were particularly low in leukemia cell lines and could not be detected by Western Blot (Fig. 4A-B, Suppl. Fig. 4C). Thus, this low FSP1 level could potentially contribute to the acute vulnerability of B-ALL cells to ferroptosis induction. To test this hypothesis, we generated B-ALL stable pools that overexpressed FSP1 (Fig. 4B) and compare their RSL3 dose-response curve with parental cells (Fig. 4C). Both pools (low or high) overexpressing FSP1 rescued sensitivity to RSL3 treatment by approximately ten-fold compared with parental cell lines, suggesting that the low endogenous constitutive levels of FSP1 contribute to the ferroptosis sensitivity in B-ALL. Furthermore, rescue levels being independent of FSP1 overexpression level indicate that weak overexpression is sufficient to inhibit ferroptosis.

**Figure 4.**
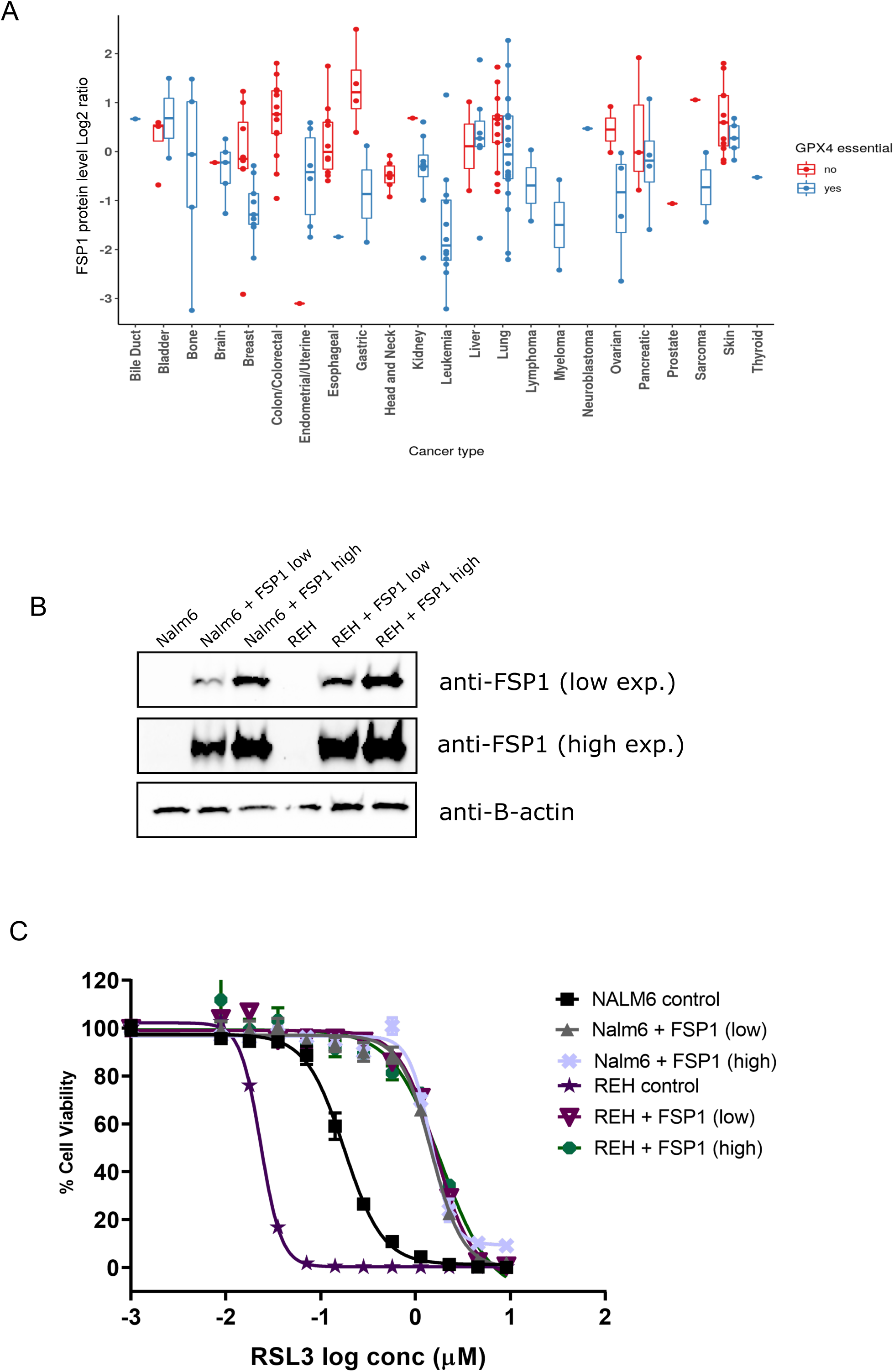
Low level of FSP1 in B-ALL contributes to ferroptosis sensitivity. A) FSP1 protein level comparison between GPX4 sensitive and non-sensitive CCLE cell lines by cancer type. (see material and method) B) FSP1 WB on parental vs FSP1 overexpressing B-ALL cells. B-actin is shown as a loading control. C) RSL3 dose-response curves in B-ALL pools overexpressing high or low levels of FPS1.

### Primary ALL PDX are also sensitive to ferroptosis induction

To validate that the sensitivity observed in B-ALL cell lines is conserved in patient-derived samples, we tested different ferroptosis inducing drugs on nine B-ALL PDX having only gone through a single round of amplification in mice previous to these tests. All xenograft samples showed high sensitivity to RSL3 treatment and were even more sensitive than the positive control cell lines, NALM6 (Fig. 5A). Furthermore, ferrostatin-1 treatment rescued RSL3 sensitivity in all PDX, indicating that ferroptosis is the major cell death mechanism in these RSL3-treated PDX. B-ALL PDX were also sensitive to three other ferroptosis-inducing drugs - erastin, FIN56 and sulfasalazine - although all PDX were not necessarily sensitive to all drugs (Fig. 5B-D). Erastin and sulfasalazine demonstrated a similar sensitivity profile consistent with these two drugs targeting the Xc^-^ transporter. In all, these experiments confirm the extreme sensitivity of B-ALL cell lines and patient-derived samples to ferroptosis induction.

**Figure 5.**
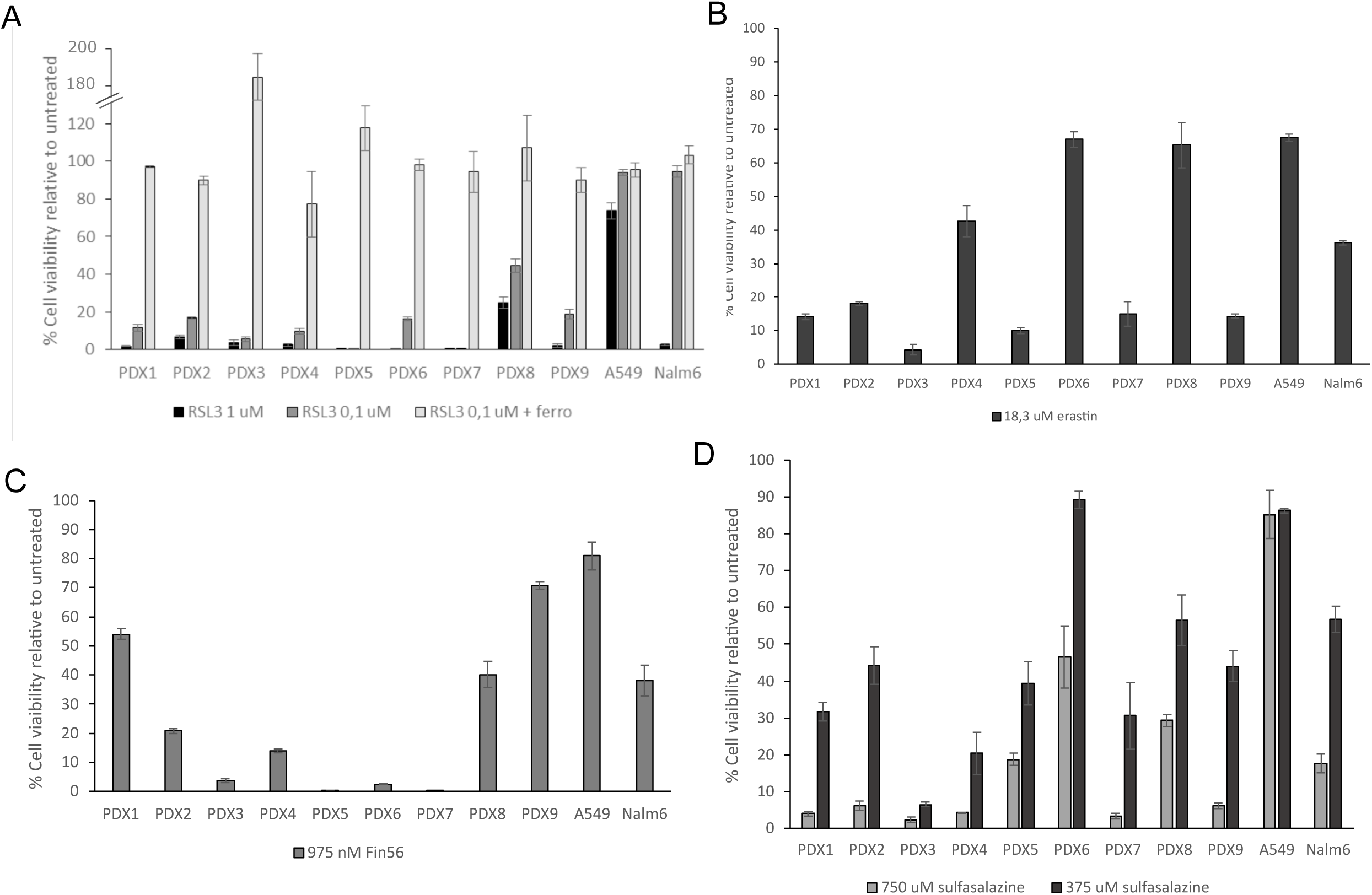
B-ALL patient derived xenografts are sensitive to ferroptosis inducing drugs. B-ALL patient-derived xenografts were treated with A) RSL3 +/- ferrostatin, B) erastin, C) Fin56, and D) sulfasalazine. Cell viability was assessed 36h post-treatment and compared to untreated xenografts.

## Discussion

Despite the continuous improvement in treatment outcome and the greater understanding of the molecular pathogenesis underlying tumor development, B-ALL still poses unresolved clinical needs, especially to relapsed patient. Developing additional therapeutic avenues by identifying new vulnerabilities is therefore critical to provide new strategies for cancer treatment. In an attempt to fulfill this gap, we performed whole-genome pooled dropout CRISPR screens in B-ALL cell lines. These screens found essential genes that were mostly shared with other cancer histotypes; these genes were enriched in core essential genes mostly implicated in general processes such as transcription, translation, proteasome, etc. In addition, these screens also identified a panel of more than three hundred genes that were essential in most ALL cell line, but not across other tumor histotypes^7^, suggesting that these genes represent unique functional vulnerabilities to B-ALL. These were enriched for well-described B-ALL specific transcription factors (*PAX5, RUNX1, TCF3, EBF1*) and signal transduction pathways (*CD79A, CD79B, PIK3CD*), but also novel vulnerabilities not previously linked to B-ALL, several of which directly regulate ferroptosis induction.

Ferroptosis is a recently described iron-dependent cell death pathway that is characterized by the excessive peroxidation of phospholipid at the cell membranes. This non-apoptotic cell death pathway has garnered increasing interest as a potential novel cancer therapy since “persister” cells rendered drug-tolerant through serial exposure to chemotherapeutic agents and cells that underwent an epithelial-mesenchymal transition, two states associated with resistance to cancer therapy, demonstrate exquisite sensitivity to ferroptosis induction^27,34^. The main gatekeeper for ferroptosis induction is GPX4, a glutathione peroxidase, which uses GSH as an obligate co-factor and possess the unique ability to detoxify hydroxyperoxides in complex lipids. Our findings add B-ALL lines to the cell lines or cell state that have been reported to be sensitive to GPX4 inhibition by either ferroptosis inducing drug^12,26,27^ or direct gene knock-out^35^. To our knowledge, this is the first evidence of such a sensitivity in these cells and the first demonstration that this sensitivity can be exploited therapeutically (see below). Many genes modulating ferroptosis induction are still not labelled as such by KEGG or MSigDB enrichment tools^36–38^, especially for genes involved in the multiple metabolic pathways that are indirectly regulating GPX4 activity (Fig. 6). This explains why ferroptosis pathway, even if significantly enriched, was not ranked higher in our Wikipathway analysis (Supplemental Table 5) even though many B-ALL essential genes were involved in pathways related to GPX4 activity, such as selenocompound, lipid, mevalonate, GSH and iron metabolism (Fig. 6). We validated several of these genetic dependencies using orthogonal assays and demonstrated that inhibition of these pathways induce ferroptosis in B-ALL cells. As shown by our +/- ferrostatin-1 screens, apart from GPX4, B-ALL cells were also particularly sensitive to the depletion of genes implicated in selenocysteine synthesis (Fig. 3H). Selenocysteine incorporation into GPX4 is required for its ferroptosis inhibitor activity^32^ and co-essentiality of selenocysteine synthesis genes with GPX4 was previously demonstrated in glioblastoma cells^39^. These results also suggest that the primary role of selenocysteine metabolism in ALL cells is to synthesize GPX4 as none of the 24 other selenocysteine-containing proteins were essential in our screen. This is reminiscent of genetic deletion of selenocysteine-containing proteins in mice where KO of GPX4 is the only one that is embryonic lethal, a phenotype shared with selenocysteine tRNA KO mice^40,41^.

**Figure 6.**
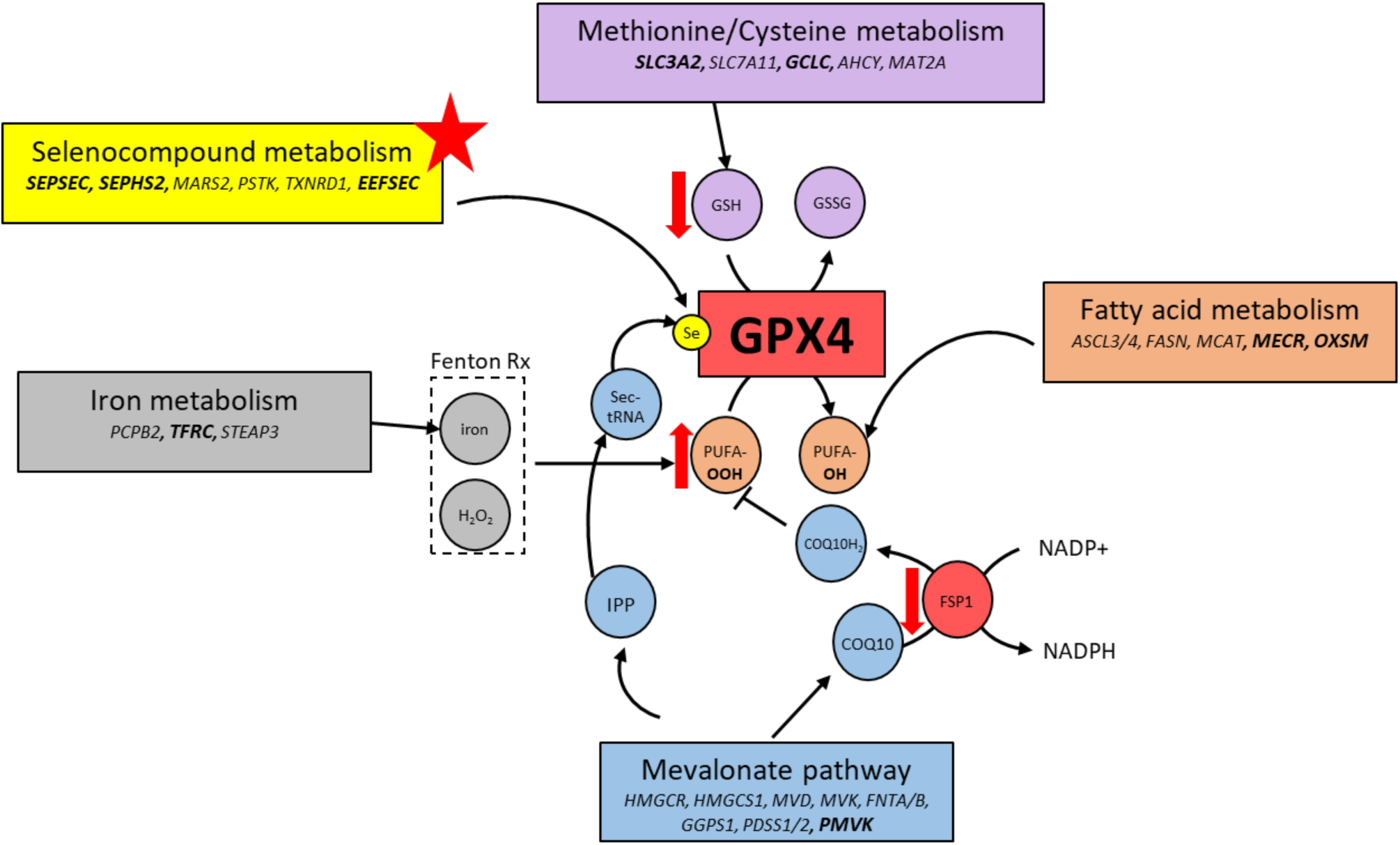
Integration of GPX4-related pathways. Several genes/pathways found essential in ALL cells potentially regulate GPX4 activity which may explain their acute vulnerability to ferroptosis induction. Genes in bold characters are found within the 319 ALL enriched essential genes.

Several elements seem to synergize to explain the high sensitivity of B-ALL cells to ferroptosis induction. High lipid ROS levels in steady-state conditions suggest that B-ALL are under constant oxidative stress. Combined with the low GSH and FSP1 antioxidant levels, it implies that these cells do not possess the buffering capacity that would normally protect them against oxidative stress. And while restoring FSP1 expression levels in B-ALL cell increases resistance to ferroptosis induction (Fig. 4C), this only partially rescues sensitivity compared to other resistant cell lines, suggesting that additional elements contribute to the extreme sensitivity of B-ALL cells to ferroptosis induction. One explanation might lie in the high levels of PUFA that were measured in B-ALL^42^ since a direct correlation between PUFA levels in cell lines and GPX4 KO sensitivity has been established by metabolite dependency association studies^43^. In addition, transcriptional repression of the pentose phosphate pathway (PPP) by B cell-specific transcriptional factor PAX5 and IKZF1 in B-ALL limits its activity and ability to cope with oxidative stress^44^. This particular vulnerability results in low levels of NADPH, which prevents GSSG reduction into GSH. The limited PPP activity appears to be controlled both by the serine-threonine protein phosphatase 2A (PP2A) and the Cyclin D3-CDK6 kinase; inhibition of the latter reduces the flow of carbon through the PPP in favor of glycolysis^45^. Cyclin D3-CDK6 kinase regulates the switch from glycolysis to the PPP by directly phosphorylating and inhibiting key enzymes, which catalyse key rate-limiting steps in the glycolysis cascade such as 6-phosphofructokinase (PFK1), pyruvate kinase M2 (PKM2), but also GPI, PGK1, ENO1, and PKM. Notably, the last four enzymes along with Cyclin D3 (CCND3) and CDK6 are all considered “B-ALL enriched essential genes” according to our screens. Overall, these results, along with the one presented in this paper, suggest that the low PPP activity found in B-ALL, which leads to low levels of NADPH and GSH, restricts their ability to cope with oxidative stress. This strenuous balance can be easily tipped toward ferroptosis induction when GPX4 is inhibited.

Most conventional chemotherapeutic agents currently use for treating ALL patients were approved more than 40 years ago and dosage and schedule have been fully optimized such that modifying the current treatment regimen will purportedly minimally improve outcome^46^. Recently, molecular targeted therapies such as BCR-ABL, JAK, mTOR, and proteasome inhibitors and immunotherapies such as monoclonal antibodies and CAR-T cells have shown great promises for improving outcomes^47^. However, the number of individuals benefiting from these novel therapies remain low for several reasons (expression of the target, resistance, toxicity, etc) such that both novel targets and therapeutics are needed to achieve greater clinical success. Ferroptosis sensitivity was seen across multiple B-ALL subtypes since several cell lines and PDX were sensitive to drugs inducing ferroptosis (Fig. 2, 3, 5 and 6). Because B-ALL are primed to undergo ferroptosis and multiple metabolic pathways modulate GPX4 activity, several therapeutic opportunities are potentially available to target this vulnerability. As shown by our results and others, ALL cells are sensitive to FDA approved drug such as sulfasalazine (Fig. 5D), BSO (Fig. 3A), and statins (Fig. 3E and ^48^). Since many chemotherapies have been shown to induce oxidative stress in cancer cells, combining these treatments with a ferroptosis inducing agent could help prevent the development of treatment-resistant tumor cells. Finally, according to gene/protein expression analyses, other leukemias, such as T-ALL and AML, also share similar expression profile for gene controlling sensitivity to ferroptosis induction and would also likely benefit from therapeutic strategies develop for ALL. Overall, our work identified a comprehensive set of genetic dependencies and molecular mechanisms sustaining tumorigenesis in B-ALL, some of which can be further exploited therapeutically.

## Methods

### Cell lines and Cell Culture

B-ALL cell lines used for this study were NALM6, HAL-01, REH, 697, RS4;11, RCH-ACV, Tanoue, SEM (DSMZ, Germany). A549 (lung carcinoma), MCF7 (breast adenocarcinoma) and NCI-H226 (lung squamous cell carcinoma) were obtained from ATCC. Cell lines were cultured in RPMI 1640 media (HyClone™) + 10% heat-inactivated FBS (Gibco™) + 2 mM glutamine (Gibco™) at 37°C in 5% CO2 incubators. HEK293SF-3F6 (ATCC) were grown in HyCell TransFx-H media (Hyclone™, Fisher Scientific) + 4 mM L-glutamine (HyClone™) + 0,1% Kolliphor (Sigma) in shaker flasks, at 120 rpm, 37°C in 5% CO2 incubators.

### B-ALL screens

B-ALL cells were infected with Cas9-2A-blast lentivirus followed by one-week selection with 5-15 µg/mL blasticidin (InvivoGen, CA). Cells were amplified and tested for Cas9 activity using the reporter assay described in Tzelepis *et al*.^14^ Pools with >75% Cas9 activity were used going forward. Cas9-expressing B-cell stable pools were transduced at 0.3 multiplicity of infection (M.O.I). with the pLKV2 whole genome human library, at 300-fold representation in triplicate (∼30 million cells per replicate). 24h post-infection, cells were centrifuged and resuspended in fresh media + 0.5-7.5 µg/mL puromycin (Gibco™). 48h post puromycin addition, cells were counted with Vi-Cell™ XR (Beckman Coulter, IN) using Trypan Blue for cell viability assessment and 30 million cells were pelleted per replicate for T0 time points. M.O.I. evaluation was done using CellTiter-Glo® (Promega, WI) and by comparing luminescence of infected cells without puromycin selection and luminescence of cells under puromycin selection. Luminescence was measured using a Synergy 2 luminometer (BioTek®) in 96-well white plates. 30 million cells per replicate were plated in culture and left for about 14 doublings with regular passage (2-4 days). After 14 doublings, 30 million cells per replicate were pelleted for Tf time points.

### Gene network analysis

Key pathways associated with ALL specific genes were determined by mapping ALL enriched essential genes onto the STRING protein database^49^. High confidence protein-protein interactions were retained (score > 0.7). We used Cytoscape and maximum clique centrality (MCC) to identify the top 100 hub genes^50,51^. For each hub (cluster), a KEGG analysis was performed to highlight key pathways in the network^38^.

### Drug treatments

Ferroptosis-modulating drugs were procured from Millipore Sigma (Oakville, ON, Canada). DMSO was from Santa Cruz Biotechnologies (Dallas, TX). IC50 curves and drug rescue were measured using CellTiter-Glo® (Promega, WI) and luminescence was measured as described above. For drug rescue experiments, both drugs were added simultaneously.

### Flow Cytometry analysis

Cells treated with 1 µM RSL3 for 4h were stained with 200 µM C11 581/591 BODIPY™ or Annexin V Pacific Blue™ conjugate (Molecular Probes®) as per manufacturer protocol. A 20h 8 µM etoposide treatment was used as a control for apoptosis induction. For each condition, approximately 30,000 cells were analyzed by flow cytometry with a BD LSRFortessa™ system (BD Biosciences, CA). Data were analyzed using the BD FACSDiva™ Software (BD Biosciences).

### Screen +/- Ferrostatin

REH and SEM Cas9 pools were screened as described above, except that each pool was screened +/- 4 µM ferrostatin in parallel. Ferrostatin and puromycin were added 24h post transfection. Sequence analysis was performed using MAGeCK Maximum Likelihood Estimation (MLE) algorithm with combined data from two cell lines and the MAGeCKFlute R package (version 1.4.3) used to highlight treatment specific genes^52^.

### Fsp1 rescue experiments

Lentiviruses expressing FSP1-myc-DDK-P2A-puro were used to infect NALM6 and REH cell lines. FSP1 expression levels were validated by WB. Following puromycin selection, pools were used in parallel with parental cells to perform RSL3 IC50 curves as described above.

### Patient-derived xenografts and drug treatments

PDX were generated as described in details in Nicolas Montpas *et al*. (manuscript in preparation). Briefly, bone marrow samples of B-ALL patients were collected at diagnosis and cells were isolated by Ficoll-Paque. 0.6-5 × 10^6^ cells were injected in the tail vein of NOD-*scid*ILR2gamma^null^ (NSG) mice and blast percentage (blast%) in the blood was monitored by monthly bleeding using human CD45, CD10 and CD19 vs CD45 murine antibodies (Biolegend, CA). Cells were expanded until blast% >1. Mice were then sacrificed, and PDX were isolated by extracting pre-B cells from mice spleen. Drug treatments were done for 36h before cell viability assessment by CellTiter-Glo® (Promega, WI). Detailed information on each PDX can be found in Suppl. Table 7.

## Acknowledgments

We thank Sonia Leclerc for performing Next-Generation Sequencing experiments. We also thank Nadine Fradet and Mylène Gosselin for their contribution to early development of the experimental designs. All authors are members of the NRC-CHUSJ Collaborative Unit for Translational Research (CUTR). We would like to acknowledge CUTR for funding part of this work. This is NRC publication # 53536.

## Author Contributions

Contribution: MEL and RM designed the study and prepared the manuscript; MEL and AMG performed the experiments; MEL, MS, AMG, JSM and RM analyzed the data; KB and EH provided patient samples; and all authors edited and approved the manuscript.

