## Supplemental material for "Genome-wide CRISPR Screens Identify Ferroptosis as a Novel Therapeutic Vulnerability in Acute Lymphoblastic Leukemia"

### Supplementary Information

### Supplemental Methods

#### Plasmid /library preparation

pLenti-Cas9-2A-Blast was a gift from Jason Moffat (Addgene plasmid # 73310). pLKV2-U6gRNA5-PGKpuro2ABPF-W human whole-genome library (Addgene # 67989), pLKV2-U6gRNA5(gGFP)-PBKBFP2AGFP-W (Addgene plasmid #67980), pKLV2-U6gRNA5(BbsI)-PGKpuro2ABFP-W (Addgene plasmid #67974) and pLKV2-U6gRNA5(empty)-PBKBFP2AGFP-W (Addgene plasmid #67978) were a gift from Kosuke Yusa. pLenti-FSP1-myc-DDK-P2A-puro was obtained from Origene (#RC204934L3). All plasmids were amplified for maxiprep at 30ºC in NEB® Stable bacteria (NEB, ON) and CIRCLEGROW™ media (MP Biomedicals™, OH) except pLKV2 library plasmids that were amplified in electrocompetent Endura™ bacteria (Lucigen, WI), at a 250-fold colony representation on LB-agar (BD) plates with carbanecillin (Bioshop Canada, ON). DNA was extracted using a proprietary in-house low-endotoxin ion exchange method. Library representation was validated using next generation sequencing of 8 pmoles library on NextSeq 500 System using a NextSeq 500 mid output Reagent kit v2, 75 cycles (Illumina, CA).

#### Lentiviral production

HEK293SF-3F6 cells were transfected at a concentration of 1x$10^6$ cells/mL with pLKV2 library, pLKV2-U6gRNA5(gGFP)-PBKBFP2AGFP-W, pLKV2-U6gRNA5(empty)-PBKBFP2AGFP-W, pLenti-FSP1-myc-DDK-P2A-puro (Origene # RC204934L3) or lenti-Cas9-2A-blast plasmid using PEIpro® (Polyplus, NY) together with packaging plasmids for lentiviral production (pMDLg-Gag/Pol, pCMV-CuO-VSVG, pRSV-Rev) at a ratio of 50 (gene of interest):25:15:10. Sodium butyrate was added at a final concentration 5mM at 16h post-transfection and supernatants were concentrated 72 hours post-transfection 200-fold on 20% sucrose cushion at 37,000g for 3hrs at 4°C.

#### Sequencing of screen samples

Genomic DNA (gDNA) extraction was performed using Blood & Cell Culture DNA Maxi Kit (Qiagen) and RNase A (Qiagen). gDNA was amplified by a two-step nested PCR reaction, first using primers CAGCGGTGCTGTCCATCTG and CCATTTGGTTAGTACCGGGC for PCR1 with the NEBNext® Ultra™ II Q5® Master Mix (NEB, ON, Canada) and by performing 20 reactions per sample each containing 2.5 µg of gDNA. A single PCR2 reaction was done similarly using 20 µL of PCR1 and with an equimolar mix of D501_F1-F8 primers and a specific D800R primer for indexing (sequences shown in Supplemental Table 1). PCR2 was purified on agarose gel using Gel extraction kit (Qiagen) followed by SPRIselect (Beckman Coulter, IN) bead purification. Purified fragments were quantified using NEBNext® Library Quant Kit for Illumina® (NEB, ON, Canada) and 14.3 fmoles of each PCR2 was multiplexed for sequencing on a NextSeq 500 system using NSQ 500 v2 75 cycle Hi Output kit (Illumina, CA).

**Sequencing data processing and statistical analyses**

Reads from Illumina sequencing were trimmed at both ends with the cutadapt package to extract the 20 bases sgRNA sequences and MAGeCK was used to count and normalize the reads[1]. Counts were corrected for copy number with CRISPRcleanR[2]. Guides targeting genes expressed below 0.5 TPM based on our RNAseq results or with average guide count below 30 at the initial time point were discarded. Finally, MAGeCK was used to evaluate gene essentiality for each cell lines separately with the Robust Rank Analysis (RRA) algorithm, choosing the best p-value from the positive and negative test. ROC curves were generated using CRISPRcleanR package using BAGEL essential and BAGEL non-essential gene sets as true positive and true negative reference, respectively[2]. Principal component analysis (PCA) was done with the prcomp function of the R stats package (version 3.6.2)[3]. We identified ALL-enriched genes by comparing our significant gene list (present in >4/7 cell lines, 5% FDR) to the Broad and Sanger essential gene lists[4]. Wikipathway analysis of gene and pathway enrichment was performed using Molecular Signatures Database (MSigDB) investigation tool[5,6].

**CCLE RNAseq and proteomic data analysis**

Data from the Broad Institute Cancer Cell Line Encyclopedia (CCLE) version 19Q4 was downloaded from the depmap portal (https://depmap.org/portal/download/). Essentiality scores were used to classify cell lines into either GPX4 sensitive (< -0.7) or non-sensitive (> -0.3).

Comparisons of RNA expression or protein levels between the two groups were done using two-sided unpaired Student's t-test, whose results were corrected for multiple testing using Benjamini-Hochberg. Only proteins detected in at least 20 cell lines in each group were considered for protein level.

**Antibodies and western blotting**

Anti-GPX4 (ab125066, Abcam, ON, Canada), anti-SLC7A11 (ab175186, Abcam, ON, Canada), anti-β-actin (sc-47778, Santa Cruz Biotechnologies, Dallas, TX), anti-GAPDH (G8795, Millipore Sigma, Oakville, ON, Canada) and anti-FSP1 (sc377120, Santa Cruz Biotechnologies, Dallas, TX) antibodies were used for western blotting (WB), followed by either goat anti-rabbit IgG HRP (Santa Cruz Biotechnologies, Dallas, TX) or mIgGκ BP-HRP (sc-516102, Santa Cruz Biotechnologies, Dallas, TX) respectively. Whole cell extracts were prepared by direct lysis in 4X Laemmli buffer + DTT. SDS-PAGE 4-15% gradient gels (BioRad, Mississauga, ON) were transferred on nitrocellulose membranes (BioRad, Mississauga, ON), blocked with 5% milk (BioRad, Mississauga, ON) and the signal detected with Clarity ECL (BioRad, Mississauga, ON).

**GSH level measurements**

GSH levels from 10,000 cells/well plated the day of the experiment were measured using GSH-Glo® (Promega, WI) according to the manufacturer's recommendation. Luminescence was measured as described above.

**GPX4 KO clone generation**

GPX4 sgRNAs (#1 CACGCCCGATACGCTGAGTG and #2 CTTGGCGGAAAACTCGTGCA) were subcloned in pKLV2-U6gRNA5(BbsI)-PGKpuro2ABFP-W, using BbSI (NEB, ON, Canada) digestion and ligation. Lentiviral productions of these plasmids were done as described above. SEM and REH Cas9 stable pools were infected with GPX4#2 and GPX4#1 sgRNA lentivirus respectively and cells were immediately treated with 4 µM ferrostatin and selected with 1-7.5 µg/mL puromycin (Gibco™). Cells resistant to puromycin were plated by limiting dilution into 384-well plates and clones were grown for two weeks (with addition of ferrostatin after about a week), before being transferred in two replicates 96-well plates, one without and one with 4 µM ferrostatin, respectively. After four days, a fraction of each well (+/- ferrostatin) was measured for cell viability using CellTiter-Glo® (Promega, WI). Luminescence of cells treated with or without 4 µM ferrostatin were compared to identify clones that had completely died without ferrostatin treatment. These clones were further evaluated by Sanger Sequencing for GPX4 KO, using TCCCTGCTCAGCTTCCTTTG and GCCCTTGGGTTGGATCTTCA primers for PCR on extracted gDNA.


**Supplemental Figure 1**. A) Example of Cas9 reporter assay in SEM Cas9 stable pool using pLKV2-U6gRNA5(empty)-PBKBFP2AGFP-W and pLKV2-U6gRNA5(sgGFP)-PBKBFP2AGFP-W plasmids. GFP/BFP ratio in cells infected with sgGFP plasmid reflects Cas9 activity. B) List of screened B-ALL cell lines with the respective % Cas9 activity of derived pools C) Precision / Recall curves for screened cell lines. ROC (receiver operating characteristics) curves at sgRNA level with BAGEL essential and non-essential genes as reference. D) Principal Component analysis of screen results (T0 and Tf) for B-ALL cell lines.

**Supplemental Figure 2.** A) C11 BODIPY™ vs Annexin V staining of RSL3 treated RS4;11 cells. 8 uM Etoposide +/- ZVADK caspase inhibitor treatments are shown as positive controls. B) Lipid Ros level (C11 BODIPY™) measurement at steady-state in ALL cell lines compared to non-GPX4 sensitive (non-ALL) cell lines. ** Unpaired t-test pvalue <0,01. C) Sensitivity of ALL vs non- ALL cell lines to deferoxamine treatment. Cell viability was evaluated after 24h treatment.

**Supplemental Figure 3.** A) C11 BODIPY™ vs Annexin V staining of RS4;11 treated for 24h with ferroptosis inducing drugs. B) Scheme of +/- Ferrostatin-1 screens in $REH^{Cas9}$ and $SEM^{Cas9}$ pools. C) Selenocompound metabolism gene knock-out induces lipid peroxidation. RS4;11 cells were infected with lentiviruses encoding sgRNAs targeting the indicated selenocompound gene and lipid peroxidation was measured using C11 BODIPY™ 120h post-infection. D) Distribution of GPX4 essentiality across CCLE cell lines. Cell lines with a score <-0.7 or >-0.3 were deemed sensitive or non-sensitive to GPX4 depletion, respectively. E) Volcano plot of gene co-essential with GPX4 in screened CCLE cell lines. Selenocompound genes show high co-essentiality with GPX4. F) *** Unpaired t-test pvalue <0,0001 and * pvalue <0,05.

**Supplemental Figure 4**. A) GPX4 mRNA and B) protein levels distribution across CCLE cell lines in GPX4 sensitive and non-sensitive cells. C) FSP1 mRNA level comparison between GPX4 sensitive and non-sensitive CCLE cell lines by cancer type.

**Supplemental Table 1**. List of primers used for NGS.

| | |
|---|---|
| D501-F | AATGATACGGCGACCACCGAGATCTACAC TATAGCCT ACACTCTTTCCCTACACGACGCTCTTCCGATCT TTGTGGAAAGGACGAAACACCG |
| D501-F_1 | AATGATACGGCGACCACCGAGATCTACAC TATAGCCT ACACTCTTTCCCTACACGACGCTCTTCCGATCT C TTGTGGAAAGGACGAAACACCG |
| D501-F_2 | AATGATACGGCGACCACCGAGATCTACAC TATAGCCT ACACTCTTTCCCTACACGACGCTCTTCCGATCT GC TTGTGGAAAGGACGAAACACCG |
| D501-F_3 | AATGATACGGCGACCACCGAGATCTACAC TATAGCCT ACACTCTTTCCCTACACGACGCTCTTCCGATCT AGC TTGTGGAAAGGACGAAACACCG |
| D501-F_4 | AATGATACGGCGACCACCGAGATCTACAC TATAGCCT ACACTCTTTCCCTACACGACGCTCTTCCGATCT CAAC TTGTGGAAAGGACGAAACACCG |
| D501-F_6 | AATGATACGGCGACCACCGAGATCTACAC TATAGCCT ACACTCTTTCCCTACACGACGCTCTTCCGATCT TGCACC TTGTGGAAAGGACGAAACACCG |
| D501-F_7 | AATGATACGGCGACCACCGAGATCTACAC TATAGCCT ACACTCTTTCCCTACACGACGCTCTTCCGATCT ACGCAAC TTGTGGAAAGGACGAAACACCG |
| D501-F_8 | AATGATACGGCGACCACCGAGATCTACAC TATAGCCT ACACTCTTTCCCTACACGACGCTCTTCCGATCT GAAGACCC TTGTGGAAAGGACGAAACACCG |
| D801-R | CAAGCAGAAGACGGCATACGAGAT CGAGTAAT GTGACTGGAGTTCAGACGTGTGCTCTTCCGATCT CTAAAGCGCATGCTCCAGAC |
| D802-R | CAAGCAGAAGACGGCATACGAGAT TCTCCGGA GTGACTGGAGTTCAGACGTGTGCTCTTCCGATCT CTAAAGCGCATGCTCCAGAC |
| D803-R | CAAGCAGAAGACGGCATACGAGAT AATGAGCG GTGACTGGAGTTCAGACGTGTGCTCTTCCGATCT CTAAAGCGCATGCTCCAGAC |
| D804-R | CAAGCAGAAGACGGCATACGAGAT GGAATCTC GTGACTGGAGTTCAGACGTGTGCTCTTCCGATCT CTAAAGCGCATGCTCCAGAC |
| D805-R | CAAGCAGAAGACGGCATACGAGAT TTCTGAAT GTGACTGGAGTTCAGACGTGTGCTCTTCCGATCT CTAAAGCGCATGCTCCAGAC |
| D806-R | CAAGCAGAAGACGGCATACGAGAT ACGAATTC GTGACTGGAGTTCAGACGTGTGCTCTTCCGATCT CTAAAGCGCATGCTCCAGAC |
| D807-R | CAAGCAGAAGACGGCATACGAGAT AGCTTCAG GTGACTGGAGTTCAGACGTGTGCTCTTCCGATCT CTAAAGCGCATGCTCCAGAC |
| D808-R | CAAGCAGAAGACGGCATACGAGAT GCGCATTA GTGACTGGAGTTCAGACGTGTGCTCTTCCGATCT CTAAAGCGCATGCTCCAGAC |
| D809-R | CAAGCAGAAGACGGCATACGAGAT CATAGCCG GTGACTGGAGTTCAGACGTGTGCTCTTCCGATCT CTAAAGCGCATGCTCCAGAC |
| D810-R | CAAGCAGAAGACGGCATACGAGAT TTCGCGGA GTGACTGGAGTTCAGACGTGTGCTCTTCCGATCT CTAAAGCGCATGCTCCAGAC |
| D811-R | CAAGCAGAAGACGGCATACGAGAT GCGCGAGA GTGACTGGAGTTCAGACGTGTGCTCTTCCGATCT CTAAAGCGCATGCTCCAGAC |
| D812-R | CAAGCAGAAGACGGCATACGAGAT CTATCGCT GTGACTGGAGTTCAGACGTGTGCTCTTCCGATCT CTAAAGCGCATGCTCCAGAC |

**Supplemental Table 2.** Cas9 activity in each Cas9 stable pool based on reporter assay developed by [14]

| Cas9 pools | cas9 reporter assay (cas9 activity) |
|---|---|
| HAL-01 | 48,4% |
| tanoue | 96.6% |
| RCH-ACV | 93,6% |
| NALM6 | 80,5% |
| Jurkat | 34,4% |
| Molt4 | 4,7 % |
| TOM1 | 40,5% |
| REH | 85,5% |
| RPMI8402 | 75,3% |
| KOPN-8 | 46,8% |
| RS4;11 | 80,7% |
| 697 | 82,6% |
| molt3 | 88,0 % |
| SEM | 84,0% |

**Supplemental Table 5.** Wikipathways enriched in B-ALL CRISPR screens.

| Wikipathway | Nb. genes | qValue |
|---|---|---|
| Electron Transport Chain (OXPHOS system in mitochondria) | 11/106 | $1,8X10^{-7}$ |
| Oxidative phosphorylation | 9/62 | $2,3X10^{-7}$ |
| TCA Cycle | 6/18 | $4,8X10^{-7}$ |
| Amino Acid metabolism | 9/91 | $3,2X10^{-6}$ |
| One carbon metabolism and related pathways | 7/54 | $1,2X10^{-5}$ |
| Pentose Phosphate Metabolism | 3/7 | $4,4X10^{-4}$ |
| Ferroptosis | 4/40 | $4,7X10^{-3}$ |
| Cell Cycle | 6/122 | $6,6X10^{-3}$ |
| PI3K-Akt Signaling Pathway | 10/345 | $6,8X10^{-3}$ |
| Glutathione Metabolism | 3/22 | $8,5X10^{-3}$ |
| B Cell Receptor Signaling Pathway | 5/98 | $1,3X10^{-2}$ |

**Supplemental Table 7.** B-ALL patient-derived xenografts characteristics**.**

| PDX # | Gender, age at diagnosis (year) | Genetic characteristics | Amplification days in mice |
|---|---|---|---|
| #1 | Female, 2 | n/a | 116 |
| #2 | Female, 4 | ETV6 rearrangement, 3 RUNX1 copies, t(12;21) | 168 |
| #3 | Female, 7 | AML1 amplification (4-10 copies) | 68 |

| | | | |
|---|---|---|---|
| #4 | Female, 4 | Hyperdiploidy for chromosomes 4, 6, 8, 10 and 17. | 103 |
| #5 | Female, 3 | n/a | 62 |
| #6 | Female, 3 | n/a | 141 |
| #7 | Male, 16 | TCF3 (19p13) loss of a copy | 43 |
| #8 | Male, 3 | n/a | 69 |
| #9 | Female, 4 | n/a | 116 |

A

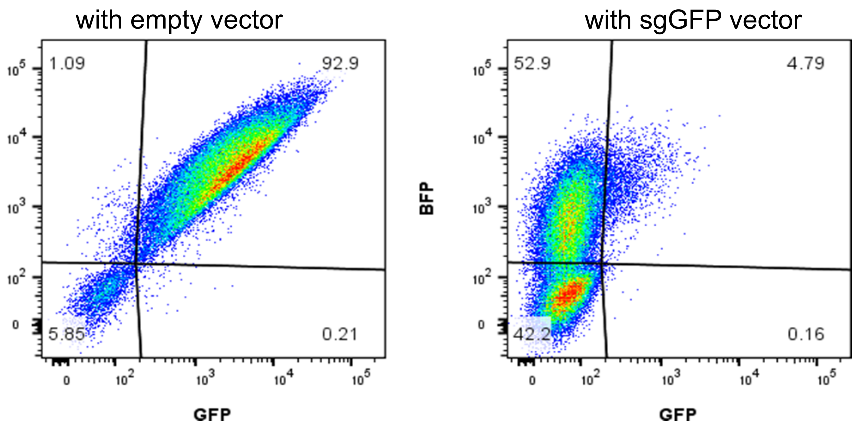


B

| Screened cell lines | % Cas9 activity of pool |
|---|---|
| Nalm6 | 80,5% |
| RS4;11 | 80,7% |
| REH | 85,5% |
| Tanoue | 96,6% |
| RCH-ACV | 93,6% |
| SEM | 84,0% |
| 697 | 82,6% |

C

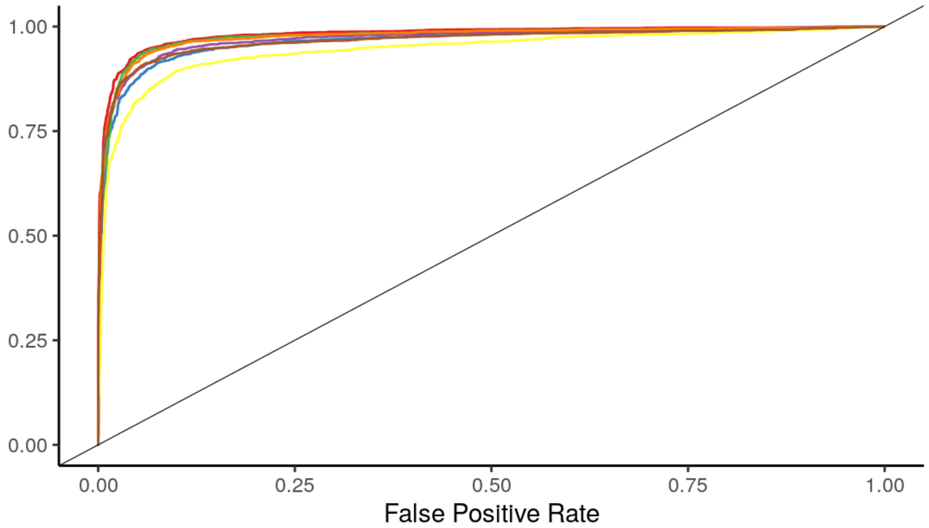


D

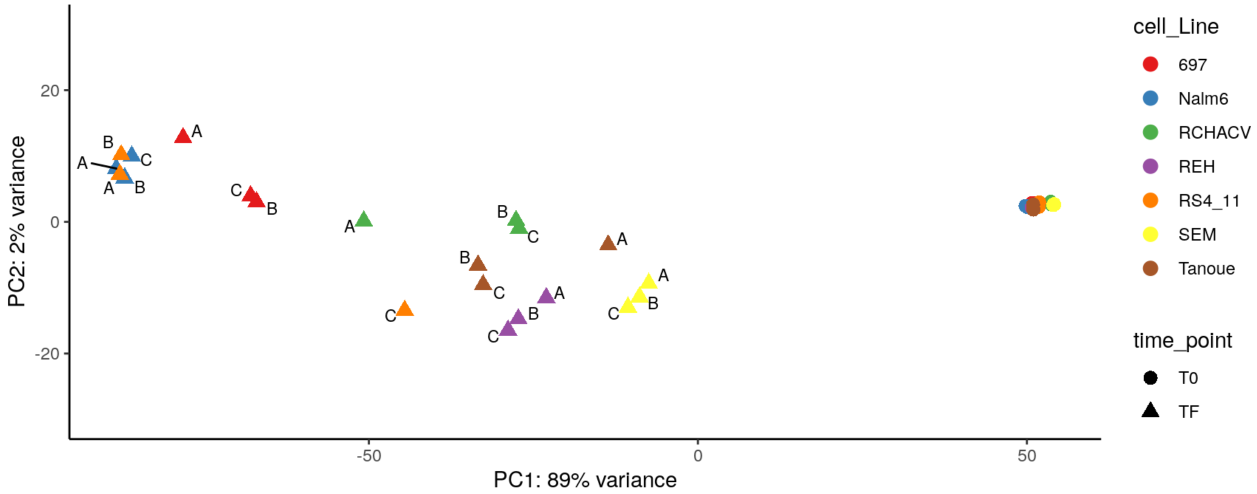


suppl. Figure 1

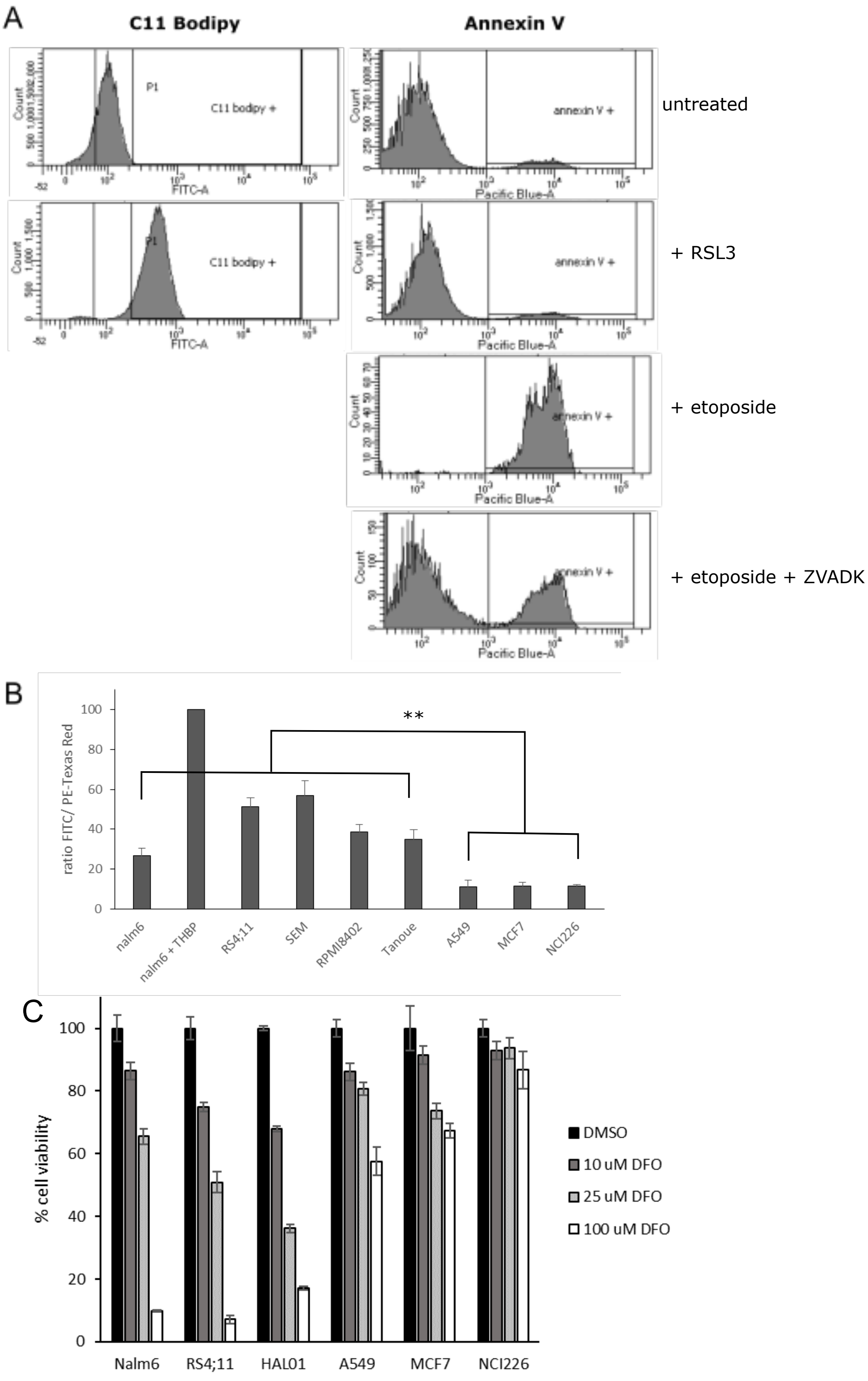


suppl. Figure 2

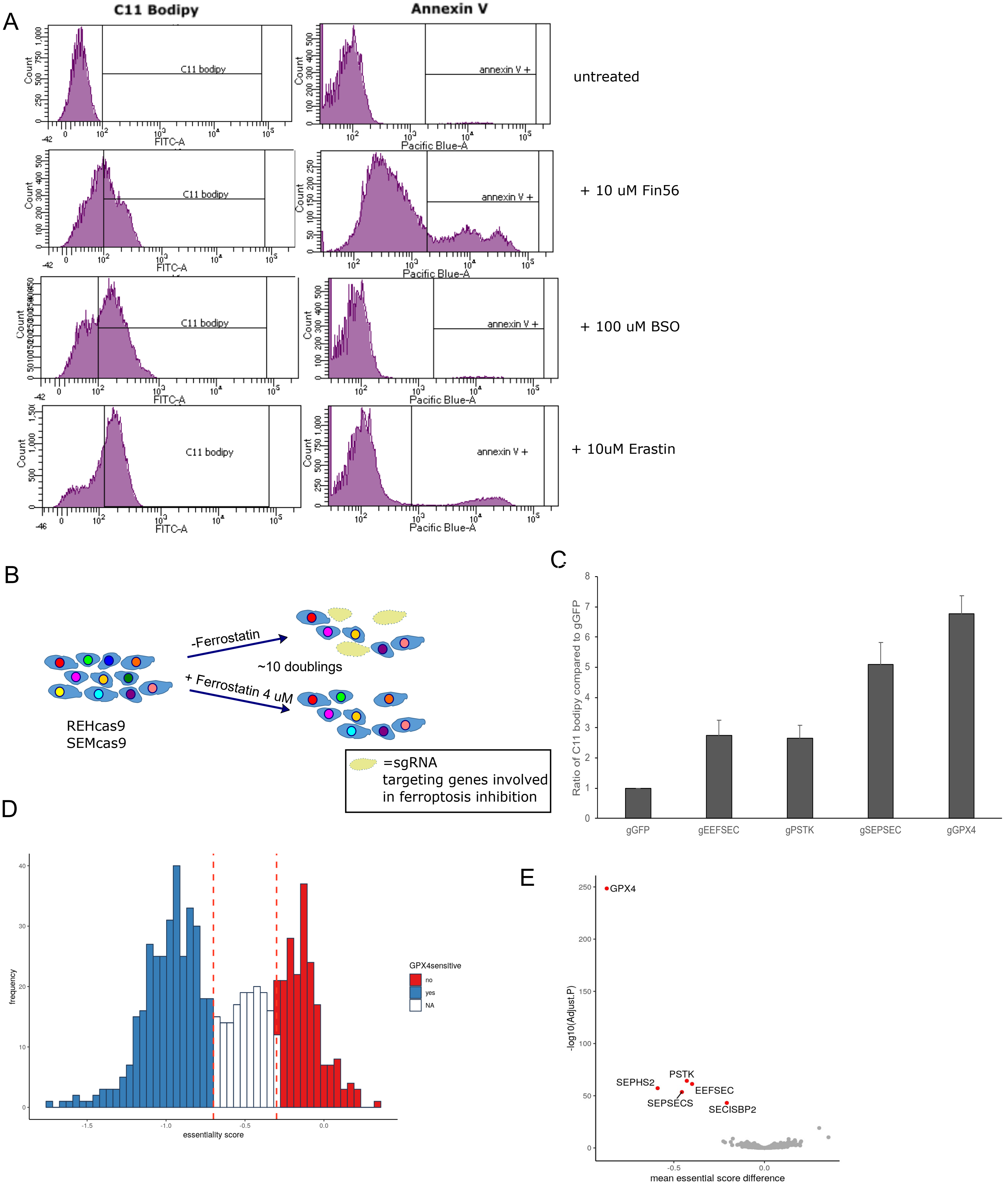


suppl. Figure 3

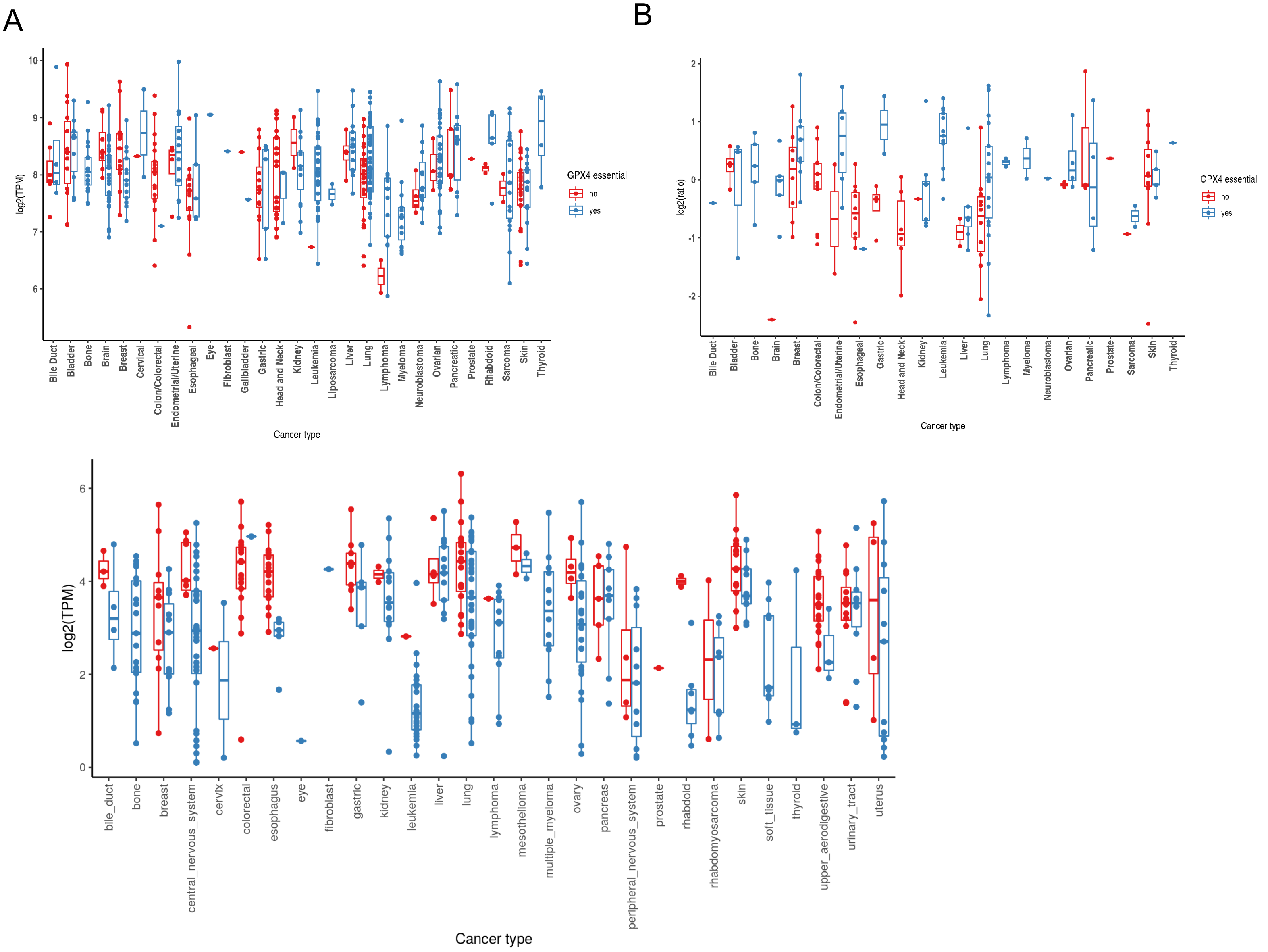


suppl. Figure 4
